# Great Ape Childhoods: Development of infant bonobos (*Pan paniscus*) and chimpanzees (*Pan troglodytes schweinfurthii*) in the wild

**DOI:** 10.1101/2025.03.14.643318

**Authors:** Jolinde M.R. Vlaeyen, Bas van Boekholt, Franziska Wegdell, Raymond Katumba, Andreas Berghänel, Martin Surbeck, Simone Pika

**Affiliations:** Institute of Cognitive Science, Comparative BioCognition, Osnabrück University, Germany; Department of Evolutionary Anthropology, University of Zurich, Zurich, Switzerland; Institute for the Interdisciplinary Study of Language Evolution, University of Zurich, Zurich, Switzerland; Sebitoli Chimpanzee Project, Great Ape Conservation Project, Fort Portal, Uganda; Department of Interdisciplinary Life Sciences, University of Veterinary Medicine, Vienna, Austria; Department of Human Evolutionary Biology, Harvard University, Cambridge, MA, USA; Department of Human Behaviour, Ecology and Culture, Max Planck Institute for Evolutionary Anthropology, Leipzig, Germany

**Keywords:** Self-Domestication Hypothesis, Extended development, Evolution, Infant, Developmental trajectories, Developmental milestones

## Abstract

Human development is marked by extended immaturity, necessitating extended care throughout infancy and childhood, facilitating advanced cognitive, social, and cultural skill acquisition. Parallels of extended development are also present in our closest living relatives, bonobos (*Pan paniscus*) and chimpanzees (*Pan troglodytes*). The Self-Domestication Hypothesis (SDH) suggests that human uniqueness stems from selection against aggression. Bonobos are also considered self-domesticated, exhibiting lower aggression and greater social tolerance, which are linked to delayed development and prolonged maternal dependence compared to chimpanzees. However, systematic, quantitative comparisons of the two species’ developmental patterns are limited and conflicting. This study addressed this gap by examining behavioural development in bonobo and chimpanzee infants aged 0-5.5 years living in two populations (Kokolopori community, Kokolopori Bonobo Reserve, DRC, N=21; Ngogo community, Kibale National Park, Uganda, N=22) in their natural environments. We specifically focused on (i) general behaviours (travel, feeding, grooming), and (ii) spatial independence. By systematically comparing developmental data and using consistent methods, we tested whether bonobo development aligns with SDH predictions. Our results showed similar developmental trajectories, with no species differences concerning ventral riding, nipple contact, or grooming. However, we found species differences regarding travel and proximity patterns, with chimpanzees exhibiting prolonged dorsal riding, bonobos travelling independently more often and maintaining greater distances from their mothers. Age, sibling presence, and maternal parity influenced behavioural patterns, but no sex differences were observed. These findings challenge assumptions of slower bonobo maturation, and highlight the importance of systematic, collaborative research on primate behavioural diversity in natural environments.

## 1. INTRODUCTION

Postnatal development is a critical phase in the life history of mammals (Purvis & Harvey, 1995), especially for altricial species such as primates. In contrast to precocial species, essential traits (e.g., motor skills) emerge gradually during early life stages instead of being fully present at birth (Harvey & Clutton-Brock, 1985). Thus, after birth, infants depend heavily on the proximity of their mothers or caretakers for comfort, security, and nourishment (Jones, 2011). Over time, they transition to greater autonomy, with mothers and caretakers gradually reducing physical contact (Maestripieri, 1995) and nursing while fostering social interactions (e.g., Pereira & Fairbanks, 2002; Pusey, 1983; van Noordwijk et al., 2009). Independence typically involves dispersal or integration into existing adult hierarchies, depending on sex and species (Jones, 2011). Together, the factors of prolonged development and maternal care have been shown to reduce mortality (Zipple et al., 2021), and support the acquisition of competencies needed to survive in complex social environments (Barton & Capellini, 2011; Maestripieri & Ross, 2004; van Noordwijk & van Schaik, 2005).

Among primates, humans show an extreme pattern of prolonged immaturity, requiring continuous care and support (Hrdy, 2001). They reach sexual maturity and adulthood only after an extended period of postnatal development (Bogin & Smith, 1996; Leigh, 2004). This prolonged development allows for the maturation of the nervous system, facilitating the acquisition of advanced cognitive, social, and cultural skills (Phillips & Shonkoff, 2000). To better understand the evolutionary factors shaping human development and potential shared patterns, comparisons with our closest living relatives, the bonobos (*Pan paniscus*) and the chimpanzees (*Pan troglodytes*), offer invaluable insights. While both species share extended juvenile periods with significant maternal support (e.g., Pereira & Fairbanks, 2002; Stanton et al., 2020; Surbeck et al., 2011; van Lawick-Goodall, 1967), their social strategies diverge in ways that likely also influence their developmental trajectories. Examining these differences will help to clarify which aspects of prolonged immaturity are uniquely human and which reflect broader evolutionary patterns among great apes.

While bonobos and chimpanzees share many developmental and behavioural traits (for a review, see Gruber & Clay, 2016), they also exhibit key differences in social dynamics that may influence their developmental trajectories. Bonobos live in predominantly female-dominated, egalitarian social systems (Furuichi, 2011; Kanō, 1992; Surbeck & Hohmann, 2013), where lethal aggressions have never been reported, neither for adults nor infants (Furuichi, 2011; Hohmann, 2001; Wilson et al., 2014). Female bonobos maintain close associations with their sons throughout life, influencing their reproductive success and social status (Furuichi, 1997; Surbeck et al., 2019; Surbeck et al., 2011; Toda et al., 2021). In contrast, chimpanzees live in hierarchical, male-dominated groups characterized by high levels of inter- and intra-group aggressions, including territorial defence, lethal aggressions (Goodall, 1986; Murray et al., 2007; Nishida, 2011), and infanticide (e.g., Arcadi & Wrangham, 1999; Watts & Mitani, 2000; Wilson et al., 2014). These differing social pressures could result in faster behavioural maturation in infant chimpanzees, as they may need to develop social and survival skills more rapidly to cope with their competitive environment, whereas the safer social conditions of bonobos may facilitate prolonged maternal dependence. For example, chimpanzee juveniles begin facing threats from adult males soon after weaning around the age of 4 to 5 years, whereas bonobos experience such threats closer to puberty around the age of 8 (Hohmann et al., 2019; Kuroda, 1989). Furthermore, chimpanzee mothers in Ngogo, Uganda, have been observed to actively support their offspring, intervening more frequently during infant attacks compared to bonobo mothers in Kokolopori, Democratic Republic of Congo (DRC), who intervene less often (Reddy et al., 2024). This difference may reflect lower aggression levels in bonobo societies and a reduced need for maternal protection. However, recent findings challenge the general view of bonobos being less aggressive: for instance, a study using focal data (Altmann, 1974) - rather than all-occurrence sampling - found that male-male agonistic interactions were significantly more frequent in one bonobo population (Kokolopori) compared to one chimpanzee population (Gombe, Tanzania; Mouginot et al., 2024). Nevertheless, crucial social differences between both species led researchers to propose that bonobos - like humans and other species (e.g., dogs; Range & Marshall-Pescini, 2022; elephants; Raviv et al., 2023) - may have undergone a process of selection known as self-domestication (Self-Domestication Hypothesis; SDH). This process is thought to have favoured traits typically associated with tameness, such as prolonged juvenile dependency, delayed behavioural maturation, and enhanced collaborative and social behaviours (e.g., Hare et al., 2012; Wrangham & Pilbeam, 2002).

Despite the assumption that bonobos exhibit a delayed behavioural maturation compared to chimpanzees, comparative research on *Pan* development is extremely limited (see Table 1). This is due to a strong bias towards studies on chimpanzees (Beck, 1982) – which benefit from extensive long-term field sites across tropical Africa (e.g., Goodall, 1986; Nakamura et al., 2015; van Lawick-Goodall, 1967; Watts, 2012) – and methodological inconsistencies across studies, as outlined in Table 1, that hinder direct comparisons. In addition, the few available studies on development provided mixed evidence, with Table 1 summarizing the existing research, underscoring variability in research settings and methodologies used. For instance, De Lathouwers and Van Elsacker (2006), comparing developmental patterns in both species living in captive settings, found differences concerning maternal proximity, grooming and nipple contact. In contrast, Lee and colleagues (2020), who studied developmental patterns in wild populations of female bonobo and chimpanzee infants and juveniles (LuiKotale, DRC, and Gombe, Tanzania), found no such differences early in development. However, after the age of 3, they reported that female bonobo infants spent more time at greater distances from their mothers compared to female chimpanzee infants. These findings contradict earlier studies suggesting greater maternal dependence in bonobos (De Lathouwers & Van Elsacker, 2006; Fröhlich et al., 2016; Koops et al., 2015; Kuroda, 1989). However, most of them focused on one specific aspect of development only rather than investigating full developmental trajectories. These discrepancies thus highlight potential differences related to study design rather than pinpointing true species-wide patterns. First, captive environments lack natural ecological pressures that shape developmental timing and social behaviour, potentially exaggerating or dampening species differences. Second, the wild study only included female bonobos, whereas the captive study included both sexes, leaving male developmental trajectories in the wild unexplored. As a result, it remains unclear whether observed differences truly reflect species-wide patterns or are artefacts of methodological and site-specific variation. The latter factors further complicate cross-species comparisons. Even within the same site, different methods can yield conflicting results. For instance, Fröhlich and colleagues (2016) relied solely on video recordings of communicative interactions, while Lee and colleagues (2020) applied 1-minute scan sampling across diverse contexts, potentially explaining discrepancies within the same population. Additionally, new methodologies, such as stable isotope analysis, have challenged previous assumptions about weaning processes. While earlier studies based on observations of visible nursing behaviours suggested similar weaning ages between bonobos and chimpanzees (e.g., Kuroda, 1989), isotope data now indicated that bonobos seem to wean 2.5–3 years later (Oelze et al., 2024). In addition, within species weaning age varies across populations (e.g., Lonsdorf et al., 2020), underscoring the role of ecological and demographic factors on behavioural output. These complexities highlight the need for caution and use of data across populations and species to tease apart whether differences reflect population or indeed species differences. Since prolonged immaturity is a hallmark of human development, understanding whether bonobos and chimpanzees differ in their development through comparative wild studies - including both sexes - can provide key insights into the evolutionary origins of extended childhood and dependency in early hominins.

**Table 1.** Overview of current research comparing bonobo and chimpanzee development across behavioural aspect, species, location (wild or captive), and the sex (if known).

| Aspect | Bonobos | Location | Sex | Chimpanzees | Location | Sex |
| --- | --- | --- | --- | --- | --- | --- |
| **Weaning** | Weaning at 4 to 5 years<sup>1</sup>; up to 7 years<sup>9</sup> | Wamba<sup>1</sup>,<br>LuiKotale<sup>9</sup> | Unspecified | Weaning at 4 to 5 years<sup>10,11</sup>; up to 7 years<sup>11</sup> | Ngogo<sup>10,11</sup>,<br>Kanywara<sup>11</sup>, Mahale<sup>11</sup>,<br>Gombe<sup>11</sup> | Both |
| **Social Proximity** | Never leave mother (3 months)<sup>1</sup> | Wamba<sup>1</sup> | Unspecified | Never leave mother (3-5 months)<sup>2,3</sup> | Gombe<sup>2</sup>, Tai<sup>3</sup> | Both |
|  | <1m away at 6 months<sup>1</sup> | Wamba<sup>1</sup> | Unspecified |  |  |  |
|  | Explore, touch siblings and unrelated individuals at 10 months<sup>1</sup> | Wamba<sup>1</sup> | Unspecified | Parting from mothers, and approach unrelated individuals at 6 months<sup>1</sup> | Unclear<sup>1</sup> | Unspecified |
|  | >10m at 3 years (but never too far to return immediately)<sup>1</sup> | Wamba<sup>1</sup> | Unspecified | Spending time out of maternal proximity at 2.5-3 years<sup>1</sup> | Unclear<sup>1</sup> | Unspecified |
|  | Spend 25% of time >5m from mother (4-5years), increasing to 50% at 5-8 years<sup>6</sup> | LuiKotale<sup>6</sup> | Females | Spending 25% of time >5m from mother (5-8 years)<sup>6</sup> | Gombe<sup>6</sup> | Females |
|  | Spend more time in close proximity to mother compared to chimpanzees<sup>4,7,8</sup> | LuiKotale<sup>7</sup>,<br>Wamba<sup>7,8</sup>,<br>Captivity<sup>4</sup> | Both | Spending more time at a greater distance from mother, compared to bonobos<sup>4</sup> | Captivity<sup>4</sup>, Tai<sup>7</sup>,<br>Kanywara<sup>7</sup>, Kalinzu<sup>8</sup> | Both |
|  | Maternal contact (50% of the time) from 1-2 years<sup>6</sup> | LuiKotale<sup>6</sup> | Females | Maternal contact (50% of the time) from 1-2 years<sup>6</sup> | Gombe<sup>6</sup> | Females |
| **Locomotion** | Ride mothers until 6 years (50% of the time until 4-5 years): no difference between ventral or dorsal<sup>6</sup> | LuiKotale<sup>6</sup> | Females | Riding on mothers until 4-5 years (50% of the time until 3 years), no difference between ventral or dorsal<sup>6</sup>. Gombe ends around 2.5 years<sup>2</sup>. | Gombe<sup>2,6</sup> | Both |
|  | Ventral clinging position in most cases (80%) at 2 years<sup>1</sup> | Wamba<sup>1</sup> | Unspecified | Ventral riding during the first 6-9 months<sup>2</sup>, shifts by 2 years<sup>5</sup> | Gombe<sup>2,5</sup> | Both |
|  | Dorsal riding becomes more common from 3 years onward, especially during long-distance travel<sup>1</sup> | Wamba<sup>1</sup> | Unspecified | Dorsal riding starts at around 7 months<sup>2,3</sup>, males transition earlier<sup>5</sup>; ends at 4-4.5 years<sup>5</sup> | Gombe<sup>2,5</sup>, Tai<sup>3</sup> | Both |
|  |  |  |  | Independent travel begins around 3 years, males started earlier<sup>5</sup> | Gombe<sup>5</sup> | Both |
| **Feeding** | Trying solid food, but not eating it < 1 year<sup>1</sup> | Wamba<sup>1</sup> | Unspecified | Eating of solid food starts around 6 months<sup>3</sup> | Tai<sup>3</sup> | Both |
|  | More nipple contact than chimpanzees from 3 years onwards<sup>4</sup> | Captivity<sup>4</sup> | Both | Continuous suckling during infancy<sup>2</sup>, Less nipple contact than bonobos from 3 years onwards<sup>4</sup> | Gombe<sup>2</sup>, Captivity<sup>4</sup> | Both |
|  | Suckling continues until 5-6 years<sup>6</sup> | LuiKotale<sup>6</sup> | Females | Suckling continues until 5-6 years<sup>6</sup> | Gombe<sup>6</sup> | Females |
| **Grooming patterns** | Social grooming observed from 2-3 years<sup>6</sup>, and more time grooming other group members compared to chimpanzees<sup>4</sup> | LuiKotale<sup>6</sup>,<br>Captivity<sup>4</sup> | Females | Social grooming starts around 1.5 years in Tai<sup>3</sup> and from 2-3 years in Gombe<sup>6</sup>; and spend less time grooming other group members compared to bonobos<sup>4</sup> | Gombe<sup>6</sup>,<br>Tai<sup>3</sup>, Captivity<sup>4</sup> | Both |
|  |  |  |  | Mother-Infant grooming observed rarely in the first few weeks, frequency increases with age<sup>2</sup> | Gombe<sup>2</sup> | Both |

Data collection methods: Observational, no described method<sup>1,2</sup>; Focal follows<sup>3,4,5,6,8,11</sup> (all day<sup>3,11</sup>; several hours – full day (chimps) and one hour (bonobo)<sup>6</sup>; 30-120min<sup>8</sup>) and Instantaneous scan sampling<sup>3,4,5,6,8</sup> (every 15sec<sup>4</sup>; 1min<sup>3,5,6</sup>; 2mins<sup>8</sup>); Video recordings of focal infants<sup>7</sup>; Faecal isotope analyses<sup>9,10</sup>. Locations: Captivity (7 Zoos across Europe), Gombe (Tanzania), Kalinzu (Uganda), Kanyawara (Uganda), LuiKotale (Democratic Republic of the Congo), Mahale (Tanzania), Ngogo (Uganda), Taï (Côte d’Ivoire), Wamba (Democratic Republic of the Congo). Sources: 1. Kuroda (1989); 2; van Lawick-Goodall (1967); 3. Bründl et al. (2021); 4. De Lathouwers and Van Elsacker (2006); 5. Lonsdorf et al. (2014) 6. Lee et al. (2020); 7. Fröhlich et al. (2016); 8. Koops et al. (2015); 9. Oelze et al. (2024); 10. Bădescu et al. (2017); 11. Lonsdorf et al. (2020).

**Table 2.** Table summarizing which data was collected as a function of observer, species and community, applied sampling methodology, duration and total observation times.

| Observer | Species and Community | Sampling Methods | Duration | Total Observation Time (hours) |
| --- | --- | --- | --- | --- |
| JV | Bonobos (FKK, EKK, KKL) | - Focal animal (continuous)<br>- Instantaneous scan (15 min intervals) | April-November 2022 | 166 |
| FW | Bonobos (FKK, EKK, KKL) | - Instantaneous scan (15 min intervals) | April-October 2022 | 56 |
| BvB | Chimpanzees (West, Central) | - Focal animal (continuous)<br>- Instantaneous scan (15 min intervals) | February- September 2021 and August 2022 -February 2023 | 1244 |
| RK | Chimpanzees (West) | - Focal animal (continuous)<br>- Instantaneous scan (15 min intervals) | October 2022 - February 2023 | 261 |

## AIM

This study revisited two key predictions of the SDH for bonobos (Hare et al., 2012), suggesting that bonobos exhibit slower behavioural maturation and prolonged maternal dependence compared to chimpanzees, reflecting differences in their social systems. To do so, we examined the following two research questions:

**1. Do bonobo and chimpanzee infants differ in their general behavioural development?** To address this question, we assessed the occurrence and frequency of specific behaviours marking changes during development in the context of travel (ventral riding, dorsal riding, independent travel), feeding (nipple contact), and grooming (infant to mother). If this prediction of the SDH holds true, we expect bonobo infants to exhibit a slower onset of independent travel, feeding, and grooming, whereas chimpanzee infants would transition more rapidly toward more independent steps in these behaviours. Alternatively, if the prediction is not supported, these results may suggest that differences in general behaviour are consistent between species or may vary more at the population than at the species level.
2. **Do bonobo and chimpanzee infants differ in spatial independence from their mothers over time?** To examine this question, we assessed the occurrence and frequency of spatial proximities (<1m, 1-5m, >5m) between infants and their mothers. If this prediction of the SDH holds true, we expect bonobo infants to remain in closer proximity to their mothers for a longer period during development, while chimpanzee infants will gain spatial independence earlier. Alternatively, if the results will not support the prediction, spatial independence may not consistently differ between species or may be shaped more by population-level factors than by species-wide traits. However, this pattern must be interpreted as a developmental process rather than a consequence of social influences, such as infanticide risk.

By employing consistent, quantitative methodologies across the two species living in two different populations under active selection pressures, our study contributes to a more nuanced understanding of how social systems and ecological pressures shape development, possibly offering broader implications for the evolutionary origins of prolonged human development.

## 2. METHODS

### a. Study communities

We conducted our study at the Kokolopori Bonobo Reserve (DRC) and the Ngogo Chimpanzee Project (Kibale National Park, Uganda) from April to November 2022, and February 2021 to February 2023, respectively. Five fully habituated communities were observed: three bonobo communities (Fekako, FKK; Ekalakala, EKK; and Kokoalongo, KKL) of the Kokolopori population, and two chimpanzee communities (West and Central) of the Ngogo population. The bonobos have been monitored since 2007 through a collaboration between NGOs Vie Sauvage and Bonobo Conservation Initiative, with M.S. initiating long-term detailed research on this population in 2016 (Surbeck et al., 2017). Similarly, the chimpanzees have been studied since 1995 (Wood et al., 2017), allowing individual identification and continuous long-term data collection in both populations. For this study, we focused on infants aged 0-5.5 years (N=43; 21 males, 22 females; Table S1). For a detailed overview regarding studied infants and community sizes, see supplementary materials, S1.

### b. Behavioural Observations

#### i. Bonobo data collection

Data were collected by JV (April-November 2022) and FW (April-October 2022), applying two methodologies. JV conducted continuous focal animal sampling (Altmann, 1974; Martin & Bateson, 1993) during two-hour focal follows using Cybertracker software (v3.520) on a smartphone (Cyrus CS24) to record general behavioural data on development (see behavioural parameters below). Information such as the start and end of each focal follow, and periods when the focal animal was out of sight were also recorded. This method was complemented with instantaneous scan sampling (Altmann, 1974; Martin & Bateson, 1993) at 15-minute intervals, recording specific behaviours of the focal individual, the mother, and proximity to other individuals and the mother. FW conducted scan sampling at 15-minute intervals during one-hour follows using voice notes. JV and FW maintained a record of the total duration for which a focal had been observed per month, and gave priority to those who had been sampled less in cases where multiple focal individuals were present. Following each focal follow, a new focal subject was chosen, given consistent grouping of bonobos.

Concurrent observations were occasionally conducted, with efforts made to observe different focal subjects to minimize overlap and with overlapping scans being excluded from analyses.

Party composition data were collected every 30 minutes by local field assistants, cumulatively aggregating the presence of each individual observed within the past 30 minutes. Bonobos were continuously followed from when they emerged from their nests (05:30 -06:00) until approximately 15:00 daily, resulting in a total of 222 observation hours (FKK=28; EKK=67, KKL=127). To capture developmental changes over time, each focal infant was followed for approximately one hour per month (average= 66±34 minutes/infant/month), except for infants being under one year of age, who were followed for 30 minutes per month (average = 34±19.5 minutes/infant/month). FW followed each infant for about 30 minutes per month (average = 37±14 minutes/infant/month), regardless of age. To compare behavioural development, the data of all infants (N=21), except one orphaned individual, were included in the subsequent analyses.

#### 2.3.2. Chimpanzee data collection

Data were collected by BvB (February–September 2021; August 2022–February 2023) and RK (October 2022–February 2023) following a protocol similar to JV’s bonobo data collection (continuous focal animal sampling complemented with 15-minute instantaneous scan sampling). Unlike bonobo observations, BvB and RK chimpanzee focal follows were conducted for full days (8:00–16:00) due to better visibility of the chimpanzees, enabling the observation of one focal individual per day. Party composition data were recorded cumulatively every hour. Concurrent observations never occurred. In total, 1,505 hours of observations were collected (West=1105, Central=400). Each infant was followed approximately six hours per month (>1y: average = 461±203 minutes/infant/month; <1y: 555±249 minutes/infant/month). To compare behavioural development, infants were included in the analysis, if sufficient data had already been collected in previous months, allowing for more reliable developmental tracking.

### c. Behavioural Data/Description of parameters

Following established ethograms (e.g., Goodall, 1986; Lonsdorf et al., 2014; Nishida et al., 1999), we utilized the proxies described below to measure (1) general behavioural development, and (2) spatial independence from mothers. Behavioural data were collected via focal animal sampling, while proximity data were obtained through scan sampling (Altmann, 1974; Martin & Bateson, 1993).

1. General Development:

A. Travel (motor development)

i. *Travel ventral* – The infant is riding ventrally, being transported as it clings to the belly of the mother, gripping hair between flexed fingers and toes. The mother can still also hold a hand on the infant’s body.
ii. *Travel dorsal* – The infant is riding dorsally, being transported as it lays or sits on the back of the mother.
iii. *Travel independently* – The infant is travelling quadrupedally on its own for at least 5m, but can still be in contact with the mother.
B. Feeding (nutritional development)

i. *Nipple Contact* – The infant takes a nipple in the mouth.
C. Grooming (social development)

i. *Grooming infant to mother* – The infant pushes the hair back on a body part of the mother and picks at the exposed skin using the hands.
2. Spatial Independence:

A. < 1m – The infant is within a mothers’ arms reach (body contact possible).
B. 1-5m – The infant is within five meters from the mother. The distance between infant and mother is a maximum of five meters and more than one meter.
C. > 5m – The infant is more than five meters away from the mother.

### d. Statistical analyses

#### i. Inter-rater reliability

Since the cooperation project began after data collection was completed, an inter-rater reliability test could not be conducted for the bonobo dataset. However, JV and FW discussed all definitions of their scan data in great detail to ensure consistency and comparability of collected data and data analysis. For the chimpanzee dataset, inter-rater reliability was assessed between BvB and RK, yielding a Cohen’s Kappa of 0.63, indicating substantial reliability (Landis & Koch, 1977).

#### ii. Development

##### Statistical models

To compare developmental trajectories between bonobo and chimpanzee infants, we ran two main models. To answer question one, we used similar models per behavioural category to assess how different behavioural aspects varied with age and between species (*Locomotion:* Model 1a-c; *Feeding:* Model 2; and *Grooming*: Model 3). To answer question two, we applied a model testing how mother-infant proximities change with increasing age (Model 4). All models were fitted in R (version 4.2.2; R core Team, 2022), using the “glmer” function of the *lme4* package (version 1.1-33; Bates et al., 2015) for Models 1 to 3, and the “clmm” function of the *ordinal* package for Model 4 (version 2022.11.16; Christensen, 2019). We determined collinearity using the “vif” function of the *car* package (Fox & Weisberg, 2019). Details about model assumptions and stability tests can be found in the supplementary materials, S2.

###### Model specificities

**Question One – “Do bonobo and chimpanzee infants differ in their general behavioural development?”.**

To analyse age- and species-based differences in travel, feeding, and grooming behaviours, we applied generalized linear mixed models (GLMM; Baayen, 2008) per behaviour: Travel modes (*Ventral riding -* Model 1a, *Dorsal riding -* Model 1b, *Independent travel* - Model 1c), Feeding modes (*Nipple Contact –* Model 2), and Grooming (Model 3). For Models 1 and 2, a binomial distribution was fitted with behaviour frequency as the response variable, represented by a two-column matrix of successes and failures per individual per observed day (Baayen, 2008). The definition of success and failure depended on the specific behaviour analysed: for travel, the behaviour of interest (ventral, dorsal, or independent travel) was coded as a success, while the other two travel modes were considered failures; for feeding, nipple contact was treated as a success, with independent feeding classified as a failure. For Model 3, a binomial distribution was fitted with infant-mother grooming occurrences (Yes, No) as the response variable.

Throughout all models, fixed effects were centred and included age (z-transformed), species (Bonobo, Chimpanzee), sex (F, M), presence of a younger sibling (Yes, No), and mother parity (Primiparous, Multiparous). For *Dorsal riding*, a quadratic age term was added due to non-linear patterns. A three-way interaction (age, species, sex) and a two-way interaction (age, sibling presence) were tested and excluded if non-significant. Random intercepts included individual, mother, community (five levels), date and individual nested in date (accounting for multiple individuals observed on the same day), with random slopes for age and mother within individual, and age, sex, sibling presence, and mother parity within community. To maintain precision in fixed effects estimates and keep the Type I error rate at 5%, we included all theoretically identifiable random slopes (Schielzeth & Forstmeier, 2009). Model fit was compared to null models, comprising only sibling presence and mother parity, using likelihood ratio tests. Collinearity was low (maximum VIF_Model1a_: 1.08; VIF_Model1b_: 2.1; VIF_Model1c_: 1.8; VIF_Model2_: 1.97; VIF_Model3_: 1.28). The sample analysed included 563 observations (42 individuals) for Model 1, 541 observations (43 individuals) for Model 2, and 463 observations (43 individuals) for Model 3.

**Question Two – “Do bonobo and chimpanzee infants differ in spatial independence from their mothers over time”.**

To examine whether mother-infant proximities varied with regard to age and species, we fitted an ordinal mixed model (cumulative logit link; Agresti & Kateri, 2017) using the measured distances to mothers (three levels: <1m, 1–5m, >5m) as the response variable. Fixed effects were centred and included age (z-transformed), species, sex, presence of a younger sibling and mother parity. Additionally, the number of scans per individual was included as an offset term to account for differences in observation effort and ensure comparability across individuals. A three-way interaction (age, species, sex) was tested but we excluded it due to being non-significant. Random intercepts included individual, mother, community (five levels), date, and individual nested within date (accounting for multiple individuals observed on the same day). We also included the number of individuals and siblings within 5m to control for potential alternative interactants, excluding correlated predictors (number of males/females within 5m). Party size and daily rainfall were also included as predictors. Party size was included to account for potential influences on mother-infant proximity, as these may vary in large versus small groups, and rainfall was included as proximity decreases during rain (Vlaeyen & van Boekholt, pers. obs.). To maintain precision and control the Type I error rate (5%), we included theoretically identifiable random slopes (e.g., age within individual and mother, and age, sex, and sibling presence within community) but excluded correlations among intercepts and slopes (Schielzeth & Forstmeier, 2009). Model significance was assessed by comparing the full model to a null model, comprising only sibling presence, using a likelihood ratio test. Collinearity was negligible (maximum VIF: 1.1). The dataset included 5,004 observations for 43 individuals.

## 3. RESULTS

### a. QUESTION ONE - General behavioural Patterns

#### i. TRAVEL MODES

**VENTRAL RIDING - *Model 1a.*** In both bonobos and chimpanzees, ventral riding declined with increasing age (mean±SE = -2.57 ± 0.49, P<0.001; Fig. 1A), with no significant difference between species (mean±SE = -0.49 ± 0.38, P=0.192). The factor maternal parity had for both species a significant effect (mean±SE = 0.98 ± 0.43, P=0.021), with infants of primiparous mothers showing higher rates of ventral travel. Other factors, including sex and sibling presence, had no significant effects. Overall, the test predictors had a clear effect on the model (full vs. null model: χ2 = 15.9, df = 3, P=0.001).

**Figure 1.**
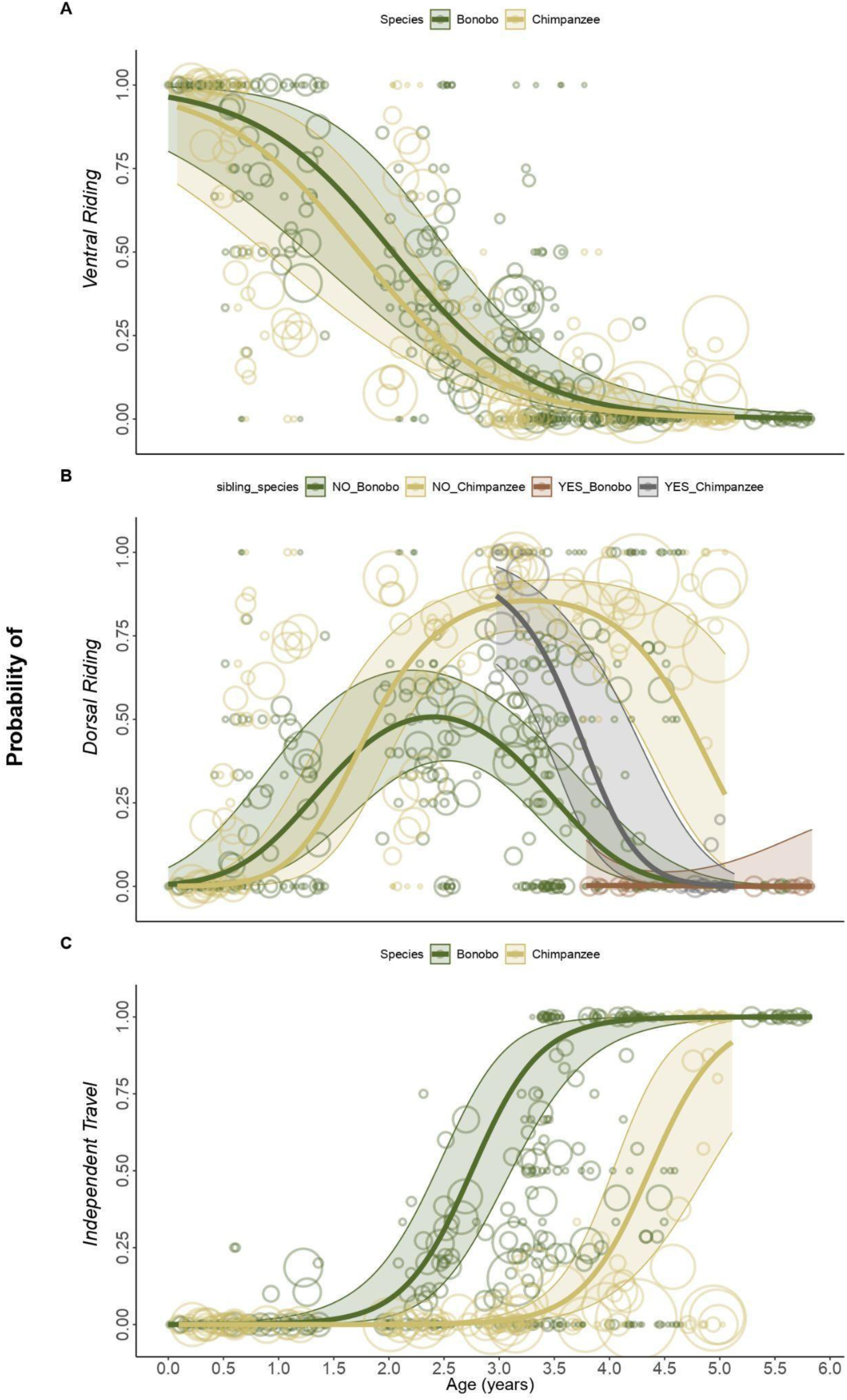
Probability figures of bonobo (dark green) and chimpanzee (yellow) infants (A) riding ventrally, (B) riding dorsally, and (C) travelling independently (the opposite as riding on mother) as a function of age (in years), including the 95% asymptotic confidence intervals. Bubbles depict the raw data with bubble area being proportional to the number of observations for each infant/age combination (range 1-60), scaled for better visibility. For figure (B), we depict the results of a different model combining species and presence of younger siblings (YES/NO) to show both statistical effects in one figure.

**DORSAL RIDING - *Model 1b.*** For both species, dorsal riding increased with age to a peak, before declining as age increased (quadratic age effect: mean±SE = -2.18 ± 0.34, P<0.001, Fig. 1B). Chimpanzee infants engaged in dorsal riding for a longer period than bonobos, as indicated by a significant interaction between age and species (mean±SE = 2.43 ± 0.69, P<0.001; Fig. 1B). For both species, infants of multiparous mothers exhibited higher dorsal riding rates (mean±SE = -1.07 ± 0.38, P=0.005), and infants without younger siblings rode dorsally longer on their mothers (negative interaction between age and sibling presence: mean±SE = -4.16 ± 1.74, P=0.017; Fig. 1B). Sex had no effect. Overall, the test predictors had a clear effect on the model (full vs null model: χ2 = 69.61, df = 7, P<0.001).

**INDEPENDENT TRAVEL - *Model 1c.*** In both bonobos and chimpanzees, independent travel increased as the infants grew older (mean±SE = 5.17 ± 0.78, P<0.001; Fig. 1C). However, bonobo infants travelled independently more frequently than chimpanzee infants, regardless of age (species effect: mean±SE = -5.09 ± 0.75, P<0.001). Visual inspection showed that this difference resulted from bonobos starting independent traveling at a younger age than chimpanzees (Fig. 1C). For both species, infants with a younger sibling also travelled independently more frequently than those without a younger sibling (mean±SE = 2.53 ± 0.78, P=0.001). Neither sex nor maternal parity had a significant effect on the test parameters. Since this model examined the inverse of riding on the mother, it also showed that chimpanzee infants travelled more often on their mothers, regardless of age. In addition, infants without younger siblings did so for longer durations. Overall, the test predictors had a clear effect on the model (full vs null model: χ2 = 29.33, df = 3, P<0.001).

Full model results and further descriptive data on travel behaviour and its developmental progression can be found in Supplementary Materials (S3).

#### ii. FEEDING MODES

**Nipple Contact - *Model 2.*** In both species, nipple contact decreased with increasing infant age (mean±SE = -1.16 ± 0.248, P<0.001; Fig. 2), with no significant difference between species (mean±SE = -0.55 ± 0.35, P=0.12). Infants without a younger sibling engaged in nipple contact for longer than those with a younger sibling. This was indicated by a significant negative interaction between sibling presence and age (mean±SE = -4.16 ± 0.96, P<0.001). Other variables, such as sex and mother parity, had no significant effects on nipple contact. Since this model examined the inverse of independent feeding, it also showed that infants with younger siblings began feeding independently earlier than those without. Overall, the test predictors had a clear effect on the model (full vs null model: χ2 = 33.3, df = 4, P<0.001).

**Figure 2.**
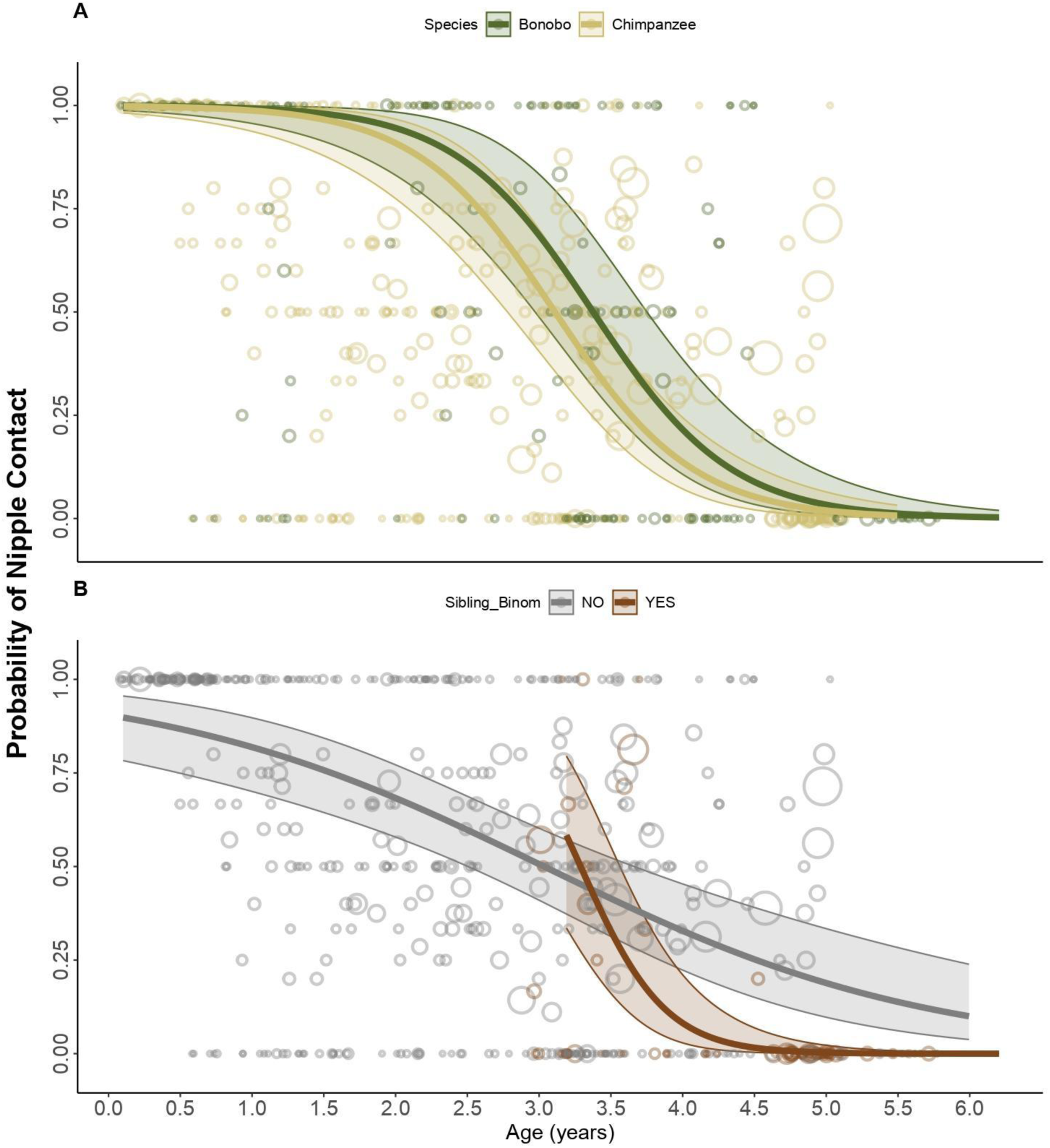
Probability of bonobo (dark green) and chimpanzee (yellow) infants nipple contact (the opposite as independent feeding) as a function of age (in years), including the 95% asymptotic confidence intervals. Bubbles depict the raw data, with bubble area being proportional to the number of observations for each infant/age combination (range 1-21), scaled for better visibility.

Full model results and further descriptive data on feeding behaviour and its developmental progression can be found in Supplementary Materials (S3).

#### iii. GROOMING

***Model 3.*** In both species, infant grooming of mothers increased with age (mean±SE = 1.11 ± 0.21, P<0.001; Fig. 3), with no significant difference between species (mean±SE = -0.29 ± 0.40, P=0.47). Other variables, such as sex, sibling presence, and mother parity, had no significant effect. Overall, the test predictors showed a clear effect on the model (full vs null model: χ² = 11.02, df = 3, P=0.011).

**Figure 3.**
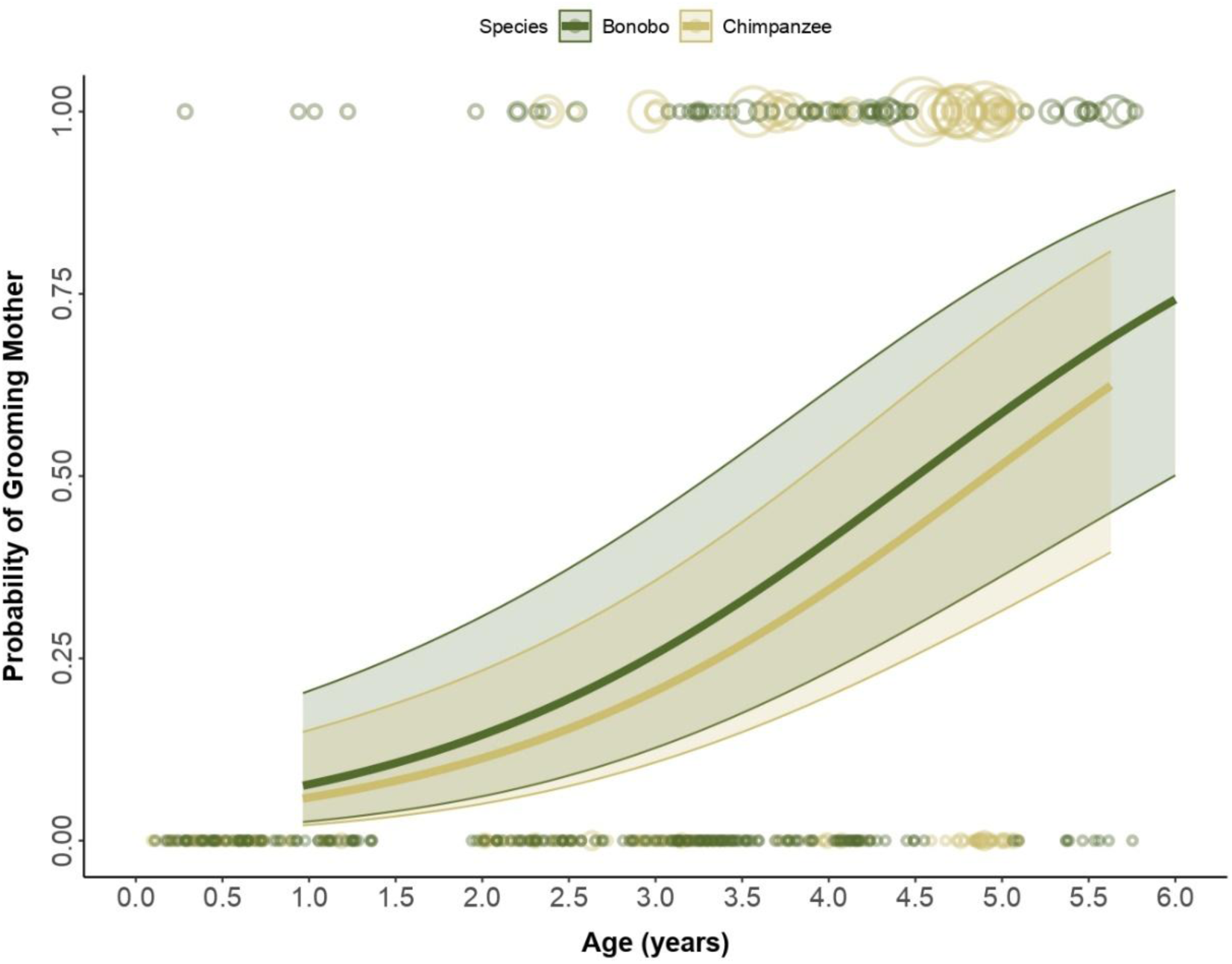
Probability of bonobo (dark green) and chimpanzee (yellow) infants grooming their mothers as a function of age (in years), including the 95% asymptotic confidence intervals. Bubbles depict the raw data with bubble area being proportional to the number of observations for each infant/age (range 0-13) combination, scaled for better visibility.

Full model results and further descriptive data on grooming behaviour and its developmental progression can be found in Supplementary Materials (S3).

### b. QUESTION TWO – Spatial independence patterns

**SPATIAL INDEPENDENCE - *Model 4.*** In both species, younger infants remained closer to their mothers than older ones, as indicated by a significant positive effect of age on spatial proximities to mothers (mean±SE = 1.46 ± 0.14, P<0.001; Fig. 4). Species differences were observed (mean±SE = -1.13 ± 0.23, P<0.001; Fig. 4), with infant bonobos being observed less frequently in the <1m category, and more frequently in the >5m category. There were no effects on sex, sibling presence, and mother parity (see Supplementary Materials S3). Overall, the model was highly significant (full vs. null model: χ² = 64.12, df = 3, P<0.001). The threshold coefficients showed that infants were about 151 times more likely to move from the ’<1m’ to ’1-5m’ proximity category (151.61 ± 0.34, P<0.001), and about 614 times more likely to move from the ’1-5m’ to ’>5m’ proximity category (614.38 ± 0.34, P<0.001). However, chimpanzee infants were 68% less likely to transition to a more distant proximity category compared to bonobos (0.32 ± 0.23, P<0.001), meaning chimpanzees remained closer to their mothers for a greater proportion of their early development phase compared to bonobos. However, there was no significant interaction between age and species, suggesting that both species transition at a similar rate, but at different ages.

**Figure 4.**
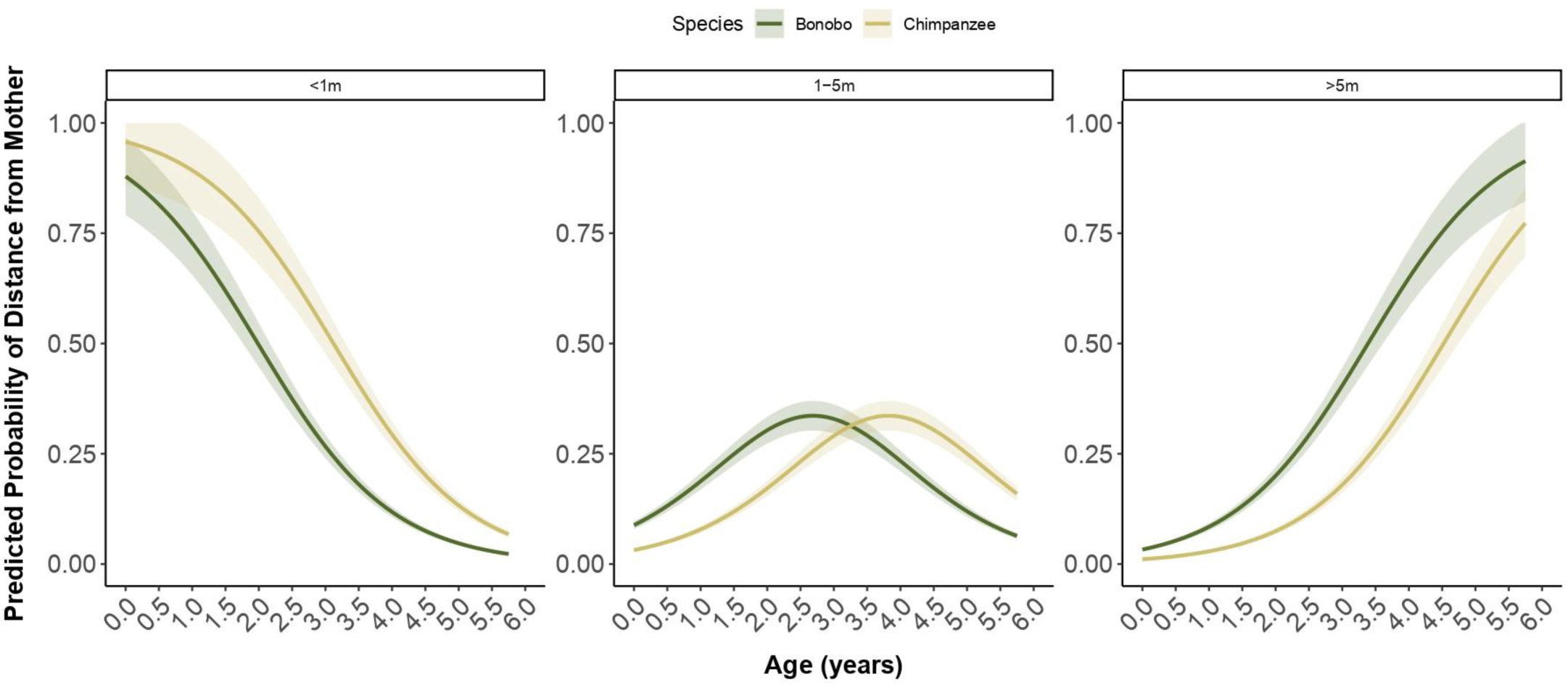
Predicted probabilities of an infant’s distance from its mother at different ages (in years) in bonobos (dark green) and chimpanzees (yellow), including the 95% asymptotic confidence intervals.

## 4. DISCUSSION

This study investigated developmental trajectories in bonobos and chimpanzees living in two populations in the wild, focusing on behavioural patterns such as travelling, feeding, grooming and spatial independence. We specifically revisited predictions of the SDH for bonobos (Hare et al., 2012; Wrangham & Pilbeam, 2002), suggesting that bonobos exhibit slower behavioural maturation and prolonged maternal dependence compared to chimpanzees. To do so, we examined the following two research questions: First, do bonobo and chimpanzee infants differ in their general behavioural development? Second, do bonobo and chimpanzee infants differ in spatial independence from their mothers over time? Overall, our results showed that age was a significant predictor across all tested parameters, validating the use of these behaviours as robust measures of development. Bonobo and chimpanzee infants followed similar developmental trajectories, with no observed differences in the behaviours of ventral riding, nursing and grooming. However, some species differences were found concerning dorsal riding, independent travel, and concerning their spatial independence. Chimpanzees rode dorsally longer than bonobos of the same age, and stayed closer to their mothers for longer than bonobos. In contrast, bonobos travelled independently more frequently at younger ages compared to infant chimpanzees. Beyond species differences, maternal parity had an effect on the way infants of both species travelled ventrally and dorsally. Infants of primiparous mothers showed higher rates of ventral travel, while those of multiparous mothers exhibited higher dorsal riding rates. In addition, the presence of a younger sibling influenced travelling and nipple contact patterns for both species. Infants without younger siblings rode dorsally for longer and nursed for a longer duration, whereas those with a younger sibling travelled independently more frequently. We did not find any effect of sex on any of the investigated behavioural parameters. In the following paragraphs, we will discuss these findings and their broader implications for development and primate evolution in more detail.

### 4.1 Development of Travelling

We identified several species-level differences for individuals of the two investigated populations regarding dorsal and independent travel. First, our findings revealed that, regardless of age, independent travel of bonobo infants was more frequent compared to chimpanzee infants, spending less time being carried by their mothers. There may be several explanations. For instance, these results may be due to species differences in locomotor behaviour. For instance, based on observations of adult individuals at Lomako, DRC, and Taï, Côte D’Ivoire, Doran (1993) suggested that bonobos are more arboreal than chimpanzees. This difference may also impact patterns of mother-infant travel, with the task of carrying an infant possibly being easier when travelling on the ground than when climbing through branches and trees. Although we do not have comparable data for chimpanzees, our data on bonobos showed that infants were carried by their mothers more often when traveling on the ground (60.5%) than when travelling in trees (39.5%). Conversely, independent travel was more common in trees (77%) compared to the ground (23%, see supplementary materials S4). These differences in locomotor behaviour may be caused by differences in habitats and structural composition. For instance, the Kokolopori Bonobo Reserve is characterized predominantly by primary, heterogeneous forests with closed canopy (e.g., Lucchesi et al., 2021; Surbeck et al., 2017) while the Ngogo landscape comprises of old-growth forests interspersed with grasslands and colonizing forest (Wing & Buss, 1970). These ecological differences may enhance and facilitate independent arboreal locomotion in bonobos at Kokolopori, in contrast to more terrestrial travel and increased reliance on maternal carrying at Ngogo. Another explanation may be that differences in travel behaviour are due to differences in the social systems, with chimpanzees facing higher risks of lethal aggressions and infanticide (Arcadi & Wrangham, 1999; Lowe et al., 2020; Watts & Mitani, 2000). Chimpanzees may thus adopt protective strategies such as prolonged carrying, as reflected by higher maternal intervention rates in offspring conflicts, compared to bonobos (Reddy et al., 2024). These differences echo protective caregiving strategies observed in human hunter-gatherers, where harsher environments necessitate extended close proximity to caregivers to enhance survival (e.g., Konner, 2017).

We also found a difference regarding dorsal riding, specifically an age-by-species difference, with older chimpanzee infants engaging in prolonged dorsal riding, compared to age-matched bonobos. These findings contrast with a study by Lee and colleagues (2020) at LuiKotale, DRC and Gombe, Tanzania, which reported prolonged dorsal riding in bonobos but not in chimpanzees. These findings may be due to population-specific ecological and social factors with the bonobo mothers frequently carrying two dependent offspring simultaneously, which rarely occurred in the chimpanzees (Lee et al., 2020). In our study populations, the chimpanzee mothers carried their infants both ventrally and dorsally concurrently, whereas such occurrences were rarely observed among the bonobos. Only three cases were observed: one post data-collection (Lily Fornof, pers. communication), and two involving the same mother—one during a stressful event (a loose dog in the forest; Vlaeyen & Wegdell, pers. obs.). Older bonobo infants occasionally attempted bipedal travel while holding onto their mothers’ backs. However, conflicts frequently arose when infants tried to ride dorsally, particularly in the presence of a new sibling (Vlaeyen, pers. obs.), suggesting mothers may actively regulate travel strategies during pregnancy or when managing multiple offspring.

Furthermore, we found high degrees of similarity between species in how maternal experience and sibling presence shaped infant travel. Infants without younger siblings rode dorsally longer and travelled independently less frequently compared to those with younger siblings. For bonobos, this pattern mirrors findings from other bonobo communities (i.e., LuiKotale) where sibling births triggered significant reductions in maternal carrying (Behringer et al., 2022). Such shifts likely reflect maternal strategies to balance the energetic demands of supporting multiple offspring while simultaneously fostering independence in older infants (Emery Thompson et al., 2016; Hrdy, 2001; Revathe et al., 2024). Additionally, maternal parity also influenced travel behaviour in both species. Infants of primiparous mothers engaged in more ventral riding, whereas infants of multiparous mothers spent more time riding dorsally. This distinction may reflect variations in maternal experience. For instance, first-time mothers, who are often less skilled, more protective and less rejective (Fairbanks, 1996; Simpson et al., 1981), may favour ventral riding for enhanced safety, while experienced mothers may encourage older infants to ride dorsally, promoting gradual independence. These findings align with other studies, including humans, showing that first-time mothers tend to invest more in their first-born (e.g., Hertwig et al., 2002; Simpson et al., 1981; Stanton et al., 2014).

In sum, our findings on infant travelling contradict prior research indicating slower maturation in bonobos during infancy and juvenility (Kuroda, 1989; Lee et al., 2020). Rather than reflecting general species differences, these results suggest population-level variation in maternal styles. For instance, mothers from both species appeared to adjust their travel strategies based on offspring age, maternal experience, and sibling presence, revealing patterns that closely parallel human caregiving (Hewlett et al., 1998; Konner, 2017). Additionally, daily travel patterns may also influence species differences in infant independence, as greater travel demands can shape maternal strategies and infant behaviour (e.g., Gruber et al., 2016). In many human hunter-gatherer societies, early infancy is characterized by near-continuous maternal carrying, with close proximity maintained through slings or body wraps (Hewlett et al., 1998). As infants grow, they transition to clinging to maternal baskets, followed by piggyback rides from caregivers by age 3 to 5, before being encouraged to walk independently after the age of five. These shifts reflect maternal strategies to balance caregiving with fostering independence, a pattern mirrored in our findings where sibling presence accelerates transitions to independent travel (Barnett et al., 2022).

### 4.2. Development of Nipple Contact

Our findings revealed no significant differences between species, as nipple contact persisted for extended periods. Furthermore, the presence of a younger sibling seemed to accelerate the weaning process, as infants with younger siblings had significantly less nipple contact for a shorter duration than infants without a younger sibling. Our findings align with research in other primate species, demonstrating that sibling birth plays a key role in shaping weaning patterns. For instance, a study on wild rhesus macaques (*Macaca mulatta*), infants that experienced weaning completion at the time of sibling birth exhibited heightened distress behaviours (Devinney et al., 2001). In contrast, weaning-related behaviours in bonobos decline gradually before sibling birth rather than ceasing abruptly (Behringer et al., 2022). This suggests that mothers may have some understanding of sibling arrival and play an active role in gradually reducing nursing opportunities. This strategy is also observed in human hunter-gatherer societies, where nipple contact often continues for up to four years but typically declines and ends during the mother’s next pregnancy, often before the birth of the next child (Konner, 2017). In the absence of a younger sibling, breastfeeding may extend to the age of 5 years (Konner, 2017), whereas in W.E.I.R.D societies, weaning tends to occur earlier due to cultural and practical constraints (e.g., Dettwyler, 2004; Runjić Babić, 2020). While data collection on confirmation of milk ingestion remains challenging in wild populations, our findings contribute to the growing evidence of prolonged maternal investment and gradual weaning in *Pan*.

### 4.3. Development of Grooming

Our results showed that the frequency of infant-mother grooming increased with infant age, supporting the limited body of research into infant-mother grooming in bonobos and chimpanzees (e.g., Nishida, 1988; van Lawick-Goodall, 1967). Similar patterns have been reported in other primate species, such as captive baboons (*Papio anubis*) and captive rhesus macaques, where grooming replaced physical carrying (Nash, 1978), thereby emphasizing the importance of tactile engagement to promote and nurture social bonds and trust (Watts, 2000). Similarly, in humans, tactile interactions – including cuddling, playful touch, and hair-brushing – fulfil comparable functions, strengthen attachment and emotion regulation, and serve as a foundation for broader social skills beyond their immediate caregivers (Phillips & Shonkoff, 2000). Although prior studies have suggested species-specific grooming dynamics, such as longer grooming sessions in subadult and adult bonobos (Girard-Buttoz et al., 2020; Sakamaki, 2013), our findings do not support these differences at the mother-infant level.

### 4.4. Spatial Independence

We found a species difference in spatial independence, with chimpanzees remaining in closer proximity to their mothers for a greater proportion of early development compared to bonobos. This was despite both species following a similar developmental trajectory. While few studies have directly compared developmental patterns of bonobos and chimpanzees, existing research generally suggested that bonobos develop more slowly and maintain closer maternal proximity for longer periods (De Lathouwers & Van Elsacker, 2006; Fröhlich et al., 2016; Koops et al., 2015). In contrast, our findings indicate the opposite pattern. They also align with results by Lee and colleagues (2020), who reported that bonobo infants spent more time at greater distances from their mothers compared to chimpanzees, albeit only beyond the age of three. In their study, bonobo female infants at LuiKotale spent 25% of their time more than 5m away from their mothers between the age of 4 to 5 years, increasing to 50% between 5 to 8 years, whereas chimpanzee female infants at Gombe did not reach the 25% threshold until the age of 5 to 8 years (Lee et al., 2020). Our results showed a comparable species difference, with bonobos observed at distances of >5m from their mothers in most scans (52.8%) by 3.5 years, increasing to 83.3% by age 5. In contrast, chimpanzee infants remained closer, with distances of >5m observed in most scans (50%) only at 4.5 years and reaching 61.7% by age 5. There may be several explanations. For instance, factors beyond developmental pace, such as exposure to danger and maternal styles, may shape proximity patterns between mothers and their infants in both bonobos and chimpanzees. Although research suggested that immature bonobos and chimpanzees experience similar levels and intensity of aggression (e.g., Furuichi, 1997; Hohmann et al., 2019; Reddy et al., 2024), a key distinction remains: while bonobos can also be aggressive toward immatures, their level of aggression has never been observed to be lethal (Hohmann et al., 2019). In contrast, chimpanzees are renowned to exhibit both infanticide and lethal aggression (Arcadi & Wrangham, 1999; Watts & Mitani, 2000; Wilson et al., 2014). This heightened risk in chimpanzees may explain why infants and mothers remain closer together. Maternal styles may also contribute to proximity differences. At Ngogo, chimpanzee mothers provided more support during aggression than bonobo mothers at Kokolopori (Reddy et al., 2024), which may explain why bonobo infants venture further while chimpanzee infants stay closer. This more protective maternal style in chimpanzees could be a response to the higher risk of lethal aggression, reinforcing maternal proximity as a protective strategy. Similarly, infant chimpanzees at Kanyawara, Uganda, moved farther from their mothers when being in parties with fewer males (Otali & Gilchrist, 2006). Our findings thus also support results from other great apes with mothers in Bornean orangutan (*Pongo pygmaeus*) actively decreasing the distance to their offspring when adult males were present (Scott et al., 2023). Another explanation may be that predator presence could influence mother-infant proximity (e.g., Förster & Cords, 2002). However, this seems unlikely as most leopard populations have been eliminated in our study populations (Watts, 2008; Surbeck, pers. Communication).

Although we found that proximity patterns differed between bonobos and chimpanzees, our results further emphasize the broader developmental pattern that infants of both species remain in close maternal proximity for an extended period. These findings highlight the prolonged dependence of the two great ape species on maternal care. Furthermore, our results align with investigations into the general social behaviour of human and nonhuman primates, with infants gradually reducing proximity to their mothers and caretakers during development (e.g., Nash, 1978). Young children of hunter-gatherer societies stay physically close (usually within 1m; Konner, 2017) to caregivers until about 4 to 5 years of age, transitioning gradually to more independent exploration while remaining under supervision (Broesch et al., 2021; Konner, 2017).

### 4.5. Future research

Our findings offer valuable insights into great ape development, by using consistent parameters across two populations and considering both sexes. Future studies could tackle several avenues to further enhance our understanding of developmental trajectories, mothering strategies and behavioural diversity in *Pan*. One important next step could be to expand research across a broader range of age classes. For instance, behaviours observed up to the age of 5 may persist beyond this age, underscoring the need for longitudinal studies covering a broader age range. Additionally, collaboration among researchers across multiple field sites and institutions have been instrumental in advancing our understanding of primate development. Large-scale collaborative data-sharing initiatives, such as 1000PAN and ManyPrimates offer promising avenues for integrating datasets and improving comparability across populations and species. Expanding the focus to include interactions of infants with siblings, peers and other adult individuals will also provide a more holistic understanding of great ape and primate social development over time. Finally, our findings emphasize the gradual and fluid nature of behavioural transitions, underscoring the importance of examining development continuously rather than solely categorizing behaviours by rigid age groups, thereby, potentially overlooking nuanced aspects of development. Moving forward, addressing discrepancies across studies through enhanced collaboration networks and data integration will be essential for building a more unified framework for primate ontogeny.

## 5. CONCLUSION

This study provides new insights into the developmental trajectories of bonobo and chimpanzee infants of two populations. Our first research question does not have a straightforward answer—while some aspects of behavioural development are conserved across species, others show clear divergence influenced by ecological and social factors. Specifically, our results indicate that bonobo and chimpanzee infants showed similarities in their development regarding ventral riding, nipple contact, and grooming, but differed with regards to dorsal and independent travel. In contrast, our second research question is conclusively answered: bonobo and chimpanzee infants differed in their spatial independence from their mothers over time. The difference in dorsal riding appears to reflect population-level variation, while species differences in independent travel and spatial proximity may be shaped by maternal strategies, environmental risk exposure, and habitat structure. These findings highlight the importance of studying multiple populations to better understand the ecological and social factors driving primate behavioural diversity. In addition, our results challenge predictions from the SDH in the form of developmental retardation for bonobos (Hare et al., 2012; Wrangham & Pilbeam, 2002) and instead suggest that, while some developmental patterns are shared across *Pan* species, species-specific differences emerge in certain behavioural domains. The observed similarities suggest shared evolutionary pressures shaping maternal strategies across great apes and humans, reflecting the adaptive trade-off between protection and fostering independence. Rather than focusing on one species as a better comparative model, these findings reinforce the need to study both species to gain a more comprehensive understanding of human ontogeny and the shared features underlying primate development. By disentangling species-specific traits from population-level variability, we can deepen our understanding of the evolutionary origins of life history traits and their relevance to human development.

## Conflict of interest statement

The authors have no conflict of interest to declare.

## Ethics statement

The present study was purely observational and non-invasive. It strictly adhered to all applicable national, and institutional guidelines for the care and use of animals, the “Animals (Scientific Procedures) Act 1986,” and the American Society of Primatologists’ “Ethical Treatment of Non-Human Primates.” Classified as a non-animal experiment under the German Animal Welfare Act (May 25, 1998, Section V, Article 7), no formal approval was required. Observers followed strict hygiene protocols, including quarantine, face masks (Kühl, 2008), and a minimum observation distance of 7m to minimize disease transmission and behavioural disruption (e.g., Köndgen et al., 2008). Permits were obtained from Harvard University, USA, Uganda Wildlife Authority (EDO-35-01), Uganda National Council for Science and Technology (NS488), Uganda, the Institut Congolais pour la Conservation de la Nature, and the Ministry of Research, DRC. Access to the Kokolopori Bonobo Reserve was granted by the communities of Bekungu, Bolamba, Yasalakose, Yetee and Yomboli.

## Author contributions

**Conceptualization:** Simone Pika; **Data curation**: Jolinde Vlaeyen, Martin Surbeck; **Formal analysis:** Jolinde Vlaeyen, Andreas Berghänel; **Funding acquisition:** Simone Pika, Martin Surbeck; **Investigation:** Jolinde Vlaeyen, Bas van Boekholt, Franziska Wegdell, Raymond Katumba; **Methodology:** Simone Pika, Jolinde Vlaeyen, Bas van Boekholt, Martin Surbeck; **Supervision, Project administration, Writing and shaping of manuscript**: Simone Pika, Martin Surbeck; **Writing - original draft:** Jolinde Vlaeyen; **Writing - review & editing:** all authors.

## Acknowledgments

We thank the people of the villages of Bekungu, Bolamba, Yasalakose, Yetee and Yomboli who granted us access to their forest, as well as all the pisteurs for their incredible help and shared knowledge of the forest. We are also very grateful to the Institut Congolais pour la Conservations de la Nature and the Ministry of Scientific Research and Technology in the DRC for their permission to work in the Kokolopori Bonobo Reserve, and the Bonobo Conservation Initiative and Vie Sauvage for support. We thank Greg Gordon, Lara Zanutto, Esa Ahmad, Chi Hsin Chen, Nicole Lahiff, Isaac Schamberg, Melissa Berthet, Morgan Rohee and Lily Fornof for their great help and support during data collection. We are very grateful to Kevin E. Langergraber, John C. Mitani, and David P. Watts for allowing us to collect data at the Ngogo Chimpanzee Project. We thank the Uganda Wildlife Authority (UWA), the Uganda National Council of Science and Technology (UNCST), and the Makerere University for permission to work at the Makerere University Biological Field Station (MUBFS). For invaluable support at the Ngogo camp and in the field, we thank Samuel Angedakin, Chris Aliganyira, Charles Birungi, Isabelle Clark, Davis Kalunga, Diana Kanweri, Brian Kamugyisha, Kevin Lee, Godfrey Mbabazi, Lawrence Ndangizi, Sebastian Ramirez Amaya, Aaron Sandel, and Alfred Tumusiime. We are also very grateful to Roger Mundry and Kayla Kolff for helpful statistical advice and discussions, and thank the whole team of the CBC for administrative support and fruitful discussions.

This research was funded by an EU-Consolidator grant (772000, TurnTaking) to SP of the European Research Council (ERC) under the European Union’s Horizon 2020 research and innovation program, and a grant by the Austrian Science Fund (FWF, P-35753 B, DOI: 10.55776/P35753) to AB. The work of BvB and RK were supported by a Leakey Foundation Research grant awarded to BvB. FW was funded by the NCCR Evolving Language, SwCSS NSF Agreement Nr.51NF40_180888.

## Notes

### Competing Interest Statement

The authors have declared no competing interest.

